# Deep Learning Outperforms Classical Machine Learning Methods in Pediatric Brain Tumor Classification through Mass Spectra

**DOI:** 10.1101/2024.01.24.577095

**Authors:** Thais Maria Santos Bezerra, Matheus Silva de Deus, Felipe Cavalaro, Denise Ribeiro, Ana Luiza Seidinger, Izilda Aparecida Cardinalli, Andreia de Melo Porcari, Luciano de Souza Queiroz, Helio Pedrini, Joao Meidanis

## Abstract

Pediatric brain tumors are the most common cause of death among all childhood cancers and surgical resection usually is the first step in disease management. During surgery, it is important to perform safe gross resection of tumors, retaining as much brain tissue as possible. Therefore, appropriate resection margin delineation is extremely relevant.

Currently available methods for tissue analysis have limited precision, are time-consuming, and often require multiple invasive procedures. Our main goal is to test whether machine learning techniques are capable of classifying the pediatric brain tissue chemical profile generated by DESI-MSI, which is mainly lipidic, into normal or abnormal tissue and into low- and high-grade malignancy subareas within each sample.

Our experiments show that deep learning methods outperform classical machine learning methods in the task of classifying brain tissue from DESI-MSI mass spectra, both in normal versus abnormal tissue, and, for malignant tissues, in low-grade versus high-grade malignancy.

Our conclusion are based on the analysis of 34,870 annotated spectra, obtained from the neoplastic and non-neoplastic microanatomical stratification of individual samples from 116 pediatric patients who underwent brain tumor surgical resection at the Boldrini Children’s Center between 2000 and 2020. Support Vector Machines, Random, Forests, and Least Absolute Shrinkage and Selection Operator (LASSO) were among the classical machine learning techniques evaluated.

## 1 Introduction

Pediatric brain tumors are the most common cause of death among all childhood cancers ([15]). Their histological classification remains difficult, due to the presence of multiple cell types. New classes are being identified as consequence of the discovery of multiple molecular pathways related to different diseases. Even presenting a high degree of heterogeneity, surgical resection usually is the first step in disease management, which often includes maximum resection margins ([20]). However, it has already been suggested that margin reduction could improve post-surgery prognostic in some cases, due to prevention of functional brain tissue loss ([21]).

One of the approaches that can improve a neurosurgeon’s decision-making process to define resection margins correlates with the new landscape of molecular classification of cancer. By leveraging molecular characterization of cancer in a time-frame compatible with the surgical setting, screening tools such as mass spectrometry (MS) allow for rapid chemical analysis in operating room environment and could ultimately help identify the nature of biological tissue being interrogated. Amongst the most widely studied platforms for this purpose is desorption electrospray ionization (DESI), which uses fast-moving solvent droplets to extract analytes from fresh tissue and propel secondary microdroplets towards the mass analyzer [8].

The subsequent challenge is to identify which chemical components could be leveraged to the task at hand. In that regard, the knowledge that tumors share a unique phenotype of elevated cell proliferation in an adverse microenvironment, which requires changes in the metabolic phenotype, such as lipid avidity and expression, could be used for cell distinction. It has been suggested that lipid phenotype could help identify and characterize tumor cells, segregating malignant tumors from their benign counterparts, as well as low- and high-grade tumors ([3]).

The present work proposes that information about this class of molecule, which could be analyzed using rapid screening tools such as DESI-MSI, could be useful for this type of discrimination in pediatric brain tumors using machine learning techniques, which are at the forefront of refined cancer diagnosis ([11]).

The work reported here is focused on the methodological comparison of deep learning and classical machine learning as tools for the bioinformatic analysis of histopathological findings of neurooncological relevance. It should be noted that each sample was divided into multiple subareas, in which the histopathological findings included promiscuous non-neoplastic (e.g., calcification, necrotic, and keratinization zones) as well as truly neoplastic lesions. Each sample was classified according to the WHO guidelines for tumors of the central nervous system (n). However, for the purpose of the current report, we have only considered the malignancy degree identified in each subarea of a particular sample (i.e., benign and grades I-IV malignant neoplasms), without distinction of specific tumor types. The evaluation of the relative accuracy of the mass spectral analysis and the conventional histopathological tumor type classification and its impact on the resection margins will be the subject of a separate report to be published elsewhere.

## 2 Literature Review

Machine learning techniques have already shown good accuracy in the analysis of pediatric brain tumors utilizing imaging data, such as diffusion weighted imaging (DWI) data, to distinguish between three different types (medulloblastoma, pylocitic astrocitoma, and ependimoma), which achieved an 85% accuracy with a Naive-Bayes classifier ([14]). Similar studies have been done using deep learning techniques to analyze pediatric posterior fossa tumors using T2-weighted MRI data, obtaining an overall accuracy of 92%. This study’s network model performance was even higher than some of the consulted pathologists ([17]). However, using imaging data for this kind of analysis in an intrasurgical context may be challenging.

Among the available techniques for tissue analysis, mass spectrometry has already shown promising results in rapid accurate tumor evaluation, such as the use of picosecond infrared laser desorption (PIRL) MS, which showed a 98.9% accuracy in molecular analysis of medulloblastoma using a 10 second PIRL-MS sampling ([21]). A combining of mass spectrometry with supervised machine learning was able to distinguish colorectal tumor from normal tissue with an overall accuracy of 98%, and to predict the presence of lymph node metastasis in primary tumour of endometrial cancer with an overall accuracy of 80% ([13]). With respect to machine learning techniques, recent studies suggest that deep learning may yield higher accuracy and a substantial gain in speed compared to classical methods such as support vector machines ([1]).

Furthermore, lipid heterogeneity has already been shown to play a role in cancer. DESI-MS techniques could highlight this characteristic, as shown in a glioblastoma study ([6]). Laser desorption/ionization mass spectrometry has been used for cancer screening and classification based on serum metabolite profiling ([23]).

## 3 Research Planning

### 3.1 Objectives

The main goal of this project is to assess the performance of machine learning techniques in the analysis of pediatric brain tissue, distinguishing between normal and abnormal findings (including tumor), using their lipidic profile data generated by DESI-MSI. A second goal is to assess this performance within the tumoral tissue findings, e.g., between low-grade tumor (grades I and II in WHO’s grading system), and high-grade tumor (grades III and IV in the same system).

### 3.2 Pathological Findings

Brain tumor mass spectra were collected from 116 pediatric patients operated in Boldrini Children’s Hospital between 2000 and 2020. Each slide yielded hundreds of individual mass spectra, one for each pixel in the resulting image, as shown in Figure 1. The slides scanned by the mass spectrometer were also analyzed by pathology specialists, who annotated unambiguous areas according to tissues findings. Table 1 shows the number of annotated mass spectra for each tissue finding. In the first part of the table we see the abnormal tissue findings observed, comprising 32,793 spectra. The additional 2,077 pixels annotated as normal findings add up to a total of 34,870 mass spectra. Notice that only part of the pixels in each sample have been annotated, namely, those belonging to unambiguos, uniform areas.

**Figure 1:**
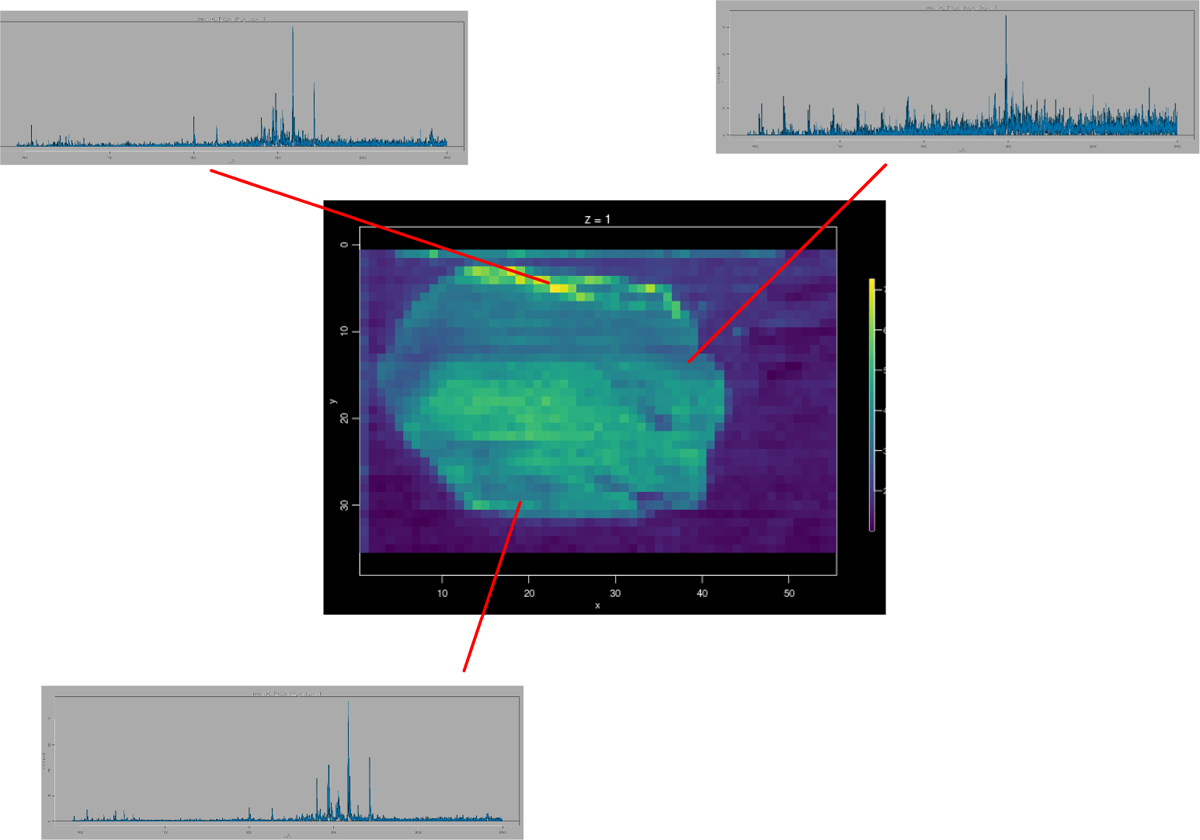
Image of scanned slide with three individual mass spectra. We have spectra for every pixel, although not every pixel has been annotated.

**Table 1:** Neoplastic and non-neoplastic findings within subareas of each pediatric brain tumor sample and the corresponding number of spectra available.

| **Tissue Finding** | **Number of Spectra** |
| --- | --- |
| Benign Neoplasm | 916 |
| Calcification | 222 |
| Keratin | 56 |
| Malignant Mesenchymal Neoplasm | 1722 |
| Metastasis | 1108 |
| Malignant Neoplasm |  |
| Grade I | 15533 |
| Grade II | 3080 |
| Grade III | 4675 |
| Grade IV | 4107 |
| Necrosis | 1374 |
| Abnormal Total | 32793 |
| Normal | 2077 |
| **Overall Total** | **34870** |

### 3.3 Machine Learning Methods

Initially, four machine learning techniques were considered: Support Vector Machine (SVM), Random Forest (RF), Multi-Layer Perceptron (MLP), and LASSO regression. The first method that was systematically explored was SVM, due to its already established performance in binary classification in Bioinformatics and its ability to deal with non-linear margins and high cardinality problems ([22]). Random Forest was considered for this project as one of the strongest tree-based models for classification problems. In general, decision trees have advantages that correlate well with our problem, given their ability to overcome the presence of redundant attributes and noise ([19]). The LASSO regression technique was considered because of its reported strong performance in very similar contexts (adult pancreatic cancer classification using spectra data) ([9]). All of those techniques were implemented using Python’s scikit-learn (sklearn) library ([16]). Deep learning approaches, such as sklearn’s MLP and Tensorflow Keras’ artificial neural networks, were considered because of a similar reason ([2]), the difference between the two being consumption of resources and flexibility, both higher in the latter.

### 3.4 Samples

The project was approved by the Institutional Review Board of Boldrini Children’s Hospital (CAAE: 38849614.2.0000.5376). The samples followed the storage flow of the institution’s Biobank (CAAE: 25574714.9.0000.5376). Only samples with consent from patients or guardians to keep the material, including later use in research projects, were selected for this project.

Tissue specimens from neurosurgical procedures were sectioned using a CM 1850 Leica cryostat (Leica® Microsystems) with a chamber at −20 °C. 14 µm thick sections were mounted onto silanized microscope glass slides (Perfecta lab) and stored at −80°C until the DESI-MSI analysis. Before being analyzed, the slides were thawed and dried by evaporation.

MS measurements were performed using a 2D Omni Spray DESI imaging platform (Prosolia Inc., Indianapolis, IN) coupled to a linear ion trap mass spectrometer (LTQ XL, Thermo Scientific). Lab-built sprayers were adapted to the commercial Omni Spray DESI imaging stages. DESI-MSI was performed on mounted tissue slides using histologically compatible solvent dimethylformamide-acetonitrile (1:1 v/v), in negative-ion mode. Solvent flow rate was 1.1 µL/min and 150 psi of N2. Data were collected as rows (spanned 0.25 mm from each other), with a rate of 248 µm/second. Ions over the mass range m/z 180–1200 were acquired in profile mode, performed with 2 microscans and injection time of 350 ms. The automatic gain control (AGC) was deactivated and relative ion intensities were shown in mass spectra.

The raw data obtained by the mass spectrometer was converted into spreadsheets generated by MSi Reader (North Carolina State University, USA) ([18], [4]), which corresponded to regions of tissue with uniform pathological findings. In addition, raw data originated from our LTQ-XL mass spectrometer, corresponding to the full slides, including annotated and nonannotated regions, were received without annotation. A database was created to store mass spectra data as blobs. Each spectrum consists of a list with 12240 floating point values, reflecting signal intensities obtained by the spectrometer, corresponding to each mass-to-charge (*m/z*) ratio, with a resolution of 1*/*12 of an inch. Even though there are only 116 patients, the total spectrum count of 34,870 is significant for machine learning analyses ([7]).

### 3.5 Design

The general established methodology for machine learning model evaluation was followed ([5]):

1. split data into train and test;
2. generate a model with train data;
3. insert test data in the model and compare the prediction to real labels.

To comply with standard terminology in the machine learning field, we will from now on refer to the pathological findings also as labels.

Given the fact that multiple spectra belong to each single patient, we made a design decision to avoid spectra from the same patient in train and test sets simultaneously. Accordingly, the chosen splitting technique has stratified folds with non-overlapping patients. Also, data was stratified along labels maintaining the same proportion as the available data displayed in Table 1.

For the classical machine learning techniques, 15-fold cross validation (CV) performed by sklearn’s function StratifiedGroupKFold was used to estimate accuracy results. In the deep learning context, this function was also used to split train and test sets, in order to achieve the desired balance. In the normal versus abnormal classification problem, train-test split followed a 4–1 ratio. A validation set with 20% of the train data was also used to monitor training evolution. In the low- versus high-grade classification problem, train-test split was 3–1. These splits are different because of the different proportions of the most frequent label in the problem set. The proportion of labels in the first classification problem, i.e., the distinction between normal and abnormal tissue findings, is 6.3% normal tissue versus 93.7% abnormal tissue. In the second classification problem, the proportion is 67.9% lowgrade findings and 32.1% high-grade findings.

#### 3.5.1 Machine Learning Model Training

To select the best SVM, RF, LASSO and MLP models for the purpose of this work, three sequential actions were performed: scale the data, select the best model hyperparameters, and build confusion matrices with the predictions of the selected model. Data was scaled used Standard Scaler from sklearn. Model selection was performed using sklearn’s Grid Search, which selected the best hyperparameters among the choices contained in a predefind parameter grid, using the best CV score. The cross-validation technique was Group-3-fold, in order to preserve time while maintaining the patient assignment restriction described previously, given the fact that Grid Search is time consuming. For a similar reason, the parameter grids prepared were constructed with the help of field specialists, which generally avoided large, time consuming grid searches. With the hyperparameters selected, the next step is to build a confusion matrix to evaluate model performance.

#### 3.5.2 Deep Learning Model Training

An incremental process of model selection was performed to achieve final results. Starting from a standard neural network architecture, several relevant parameters were tuned, including the Adam optimizer learning rate, number of epochs, and the number and size of hidden layers. Strategies such as weight initialization, data scaling, batch training, and dropout were tested to tentatively improve results. A more detailed description of this procedure is given in Section 5.1.

### 3.6 Analysis Procedure

Machine learning models were evaluated considering metrics derived from the confusion matrix, namely, accuracy, balanced accuracy, and specificity. Confusion matrices were obtained after the 15-fold CV process described in Section 3.5, in the case of classical machine learning techniques. Each matrix has the representation depicted in Table 2, where: TP stands for true positives; FP, false positives; FN, false negatives; TN, true negatives. In our application, we chose **normal findings as positive** in the normal vs. abnormal comparison, and **low-grade findings as positive** in the lowvs. high-grade tumor comparison.

**Table 2:** Confusion matrix: true positives (TP); false positives (FP); false negatives (FN); true negatives (TN).

|  | predicted positive | predicted negative |
| --- | --- | --- |
| positive | TP | FN |
| negative | FP | TN |

The quantitative metrics can be defined in terms of confusion matrix components:

- Accuracy:

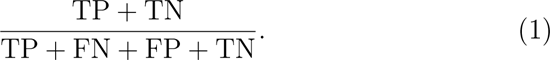

This metric is easy to calculate, and generally serves as a good comparison metric with other works.
- Balanced accuracy:

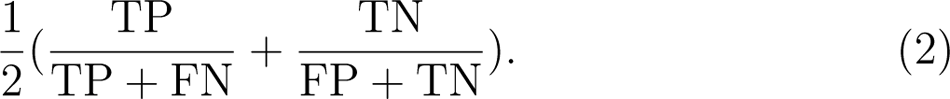

This metric was chosen as an alternative to the previous metric, especially because it considers simultaneously the true positive and the true negative rates.
- Specificity:

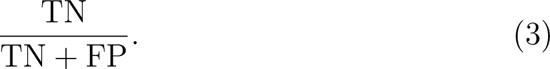

This metric was chosen because the false negatives in our problems are considered biologically more harmful than false positives. Given this fact, it is especially important to reward models that correctly classify negative spectra.

Models with the highest values of balanced accuracy and close to a diagonal confusion matrix were considered the best. We also looked at specificity, which is an important measure in our application, as explained in Section 3. Deep Learning models were evaluated using the same metrics and the computed loss of the model, which used binary cross entropy as the loss function, the most frequent option for binary classification problems.

## 4 Analysis

### 4.1 Classical Machine Learning Models

As described earlier, four machine learning techniques were considered for each classification problem: LASSO regression, SVM, RF, and MLP. The results are shown below.

#### 4.1.1 Normal versus Abnormal Distinction

We first attacked the classification problem that consisted of distinguishing between pixels with normal or abnormal label using spectra data. For each technique, we employed *Grid Search* to select the best hyperparameters within a predetermined set. In the sequel, we present the main hyperparameters, the set of values tried in the Grid Search between brackets, with the best value in boldface Hyperparameters not shown used default sklearn values.

#### SVM model

- Data scaled (standard scaler)
- Class weight = **balanced**
- gamma = [1e-3, **1e-4**, “scale”]
- Kernel = **rbf**
- C = [0.1, 1, 10, 100, **1000**]

#### Random Forest model

- Data scaled (standard scaler)
- Class weight = **balanced**
- max features = [*^√^n*, **n**] (*n* is # of features)
- min samples split = [2, 3, 4, 5, 6]
- estimators = [220, 230, 240, 250, 260, 270, 280]

#### MLP model

- alpha = [**6**, 6.5, 7, 7.5, 8]
- beta 1 = [**0.9**, 0.95, 0.8, 0.85, 0.7]
- beta 2 = [0.9999, **0.8**, 0.7]
- shuffle = [true, **false**]
- solver = **Adam**
- tolerance = [**1e-4**, 1e-3, 1e-2]
- early stopping = [**true**, false]
- epsilon = [**1e-8**, 1e-7]
- number of hidden layers = **6**
- neurons in each layer = **(40,40,40,32,32,32)**
- max iter = [250, 300, 350, 400]

#### LASSO model

- Data scaled (standard scaler)
- Class weight = **balanced**
- penalty = **L1**
- solver = **saga**
- max iter = **2000**
- C = [**0.1**, 1, 10, 50]
- tolerance = [1e-3, **1e-4**, 1e-5]

Mean confusion matrices obtained after 15-fold cross validation and the corresponding performance results can be seen in Table 3.

**Table 3:**
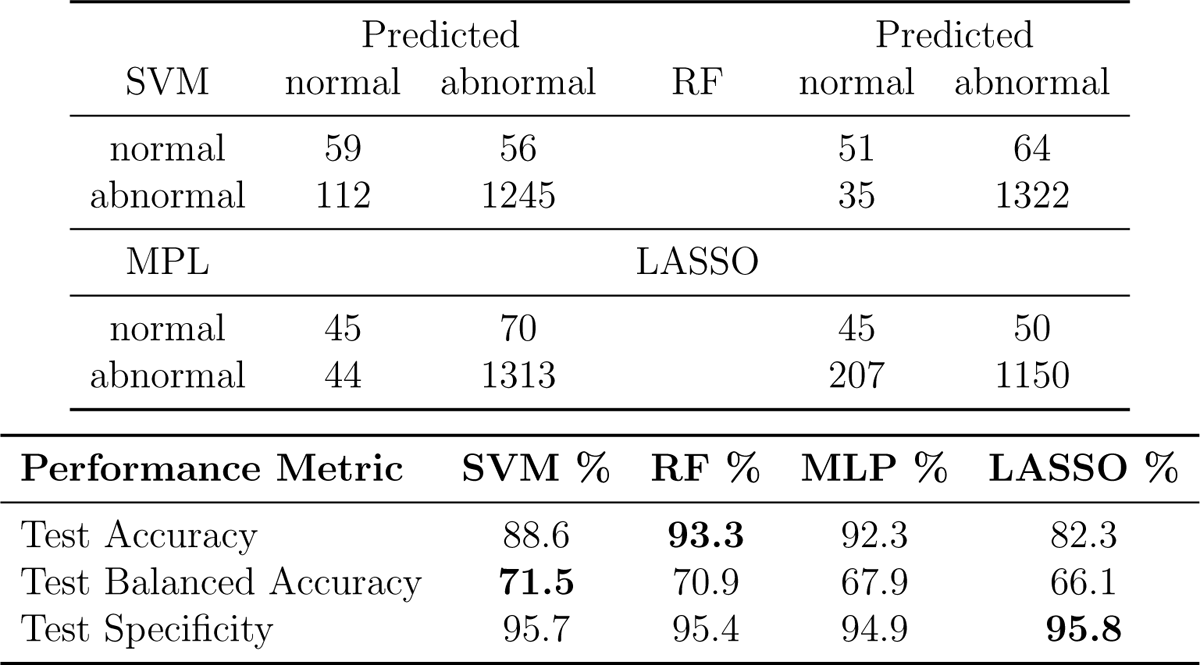
Results for classical methods in the normal versus abnormal problem: Confusion matrices and performance metrics (the best result for each row is shown in boldface).

#### 4.1.2 High grade versus low grade distinction

Given the success of the approach in normal versus abnormal classification, the same procedure was tried to distinguish between WHO low-grade and high-grade findings. Labels Grade I and II of Malignant Neoplasm are considered low-grade while labels Grade III and IV are considered high-grade. These labels were filtered out from the original dataset and relabeled as low-grade and high-grade, respectively, resulting in a dataset with 18,613 low-grade label mass spectra and 8,782 high-grade label mass spectra. Input data for the problem are unbalanced, since 67% of the total data available correspond to one label. The results were not as expressive, as can be seen in Table 4, which motivated the search for another approach.

**Table 4:** Low and high-grade classification performance results with ML models.

| Model | Test Accuracy % |
| --- | --- |
| SVM | 68.6 |
| Random Forest | 67.8 |
| MLP | 61.0 |
| LASSO | 67.7 |

### 4.2 Deep Learning Models

#### 4.2.1 High grade versus low grade

Given the poor results with regular ML techniques, a novel approach using deep learning strategies was conducted. The neural network architecture that achieved the best results had the following parameters:

6 hidden layers with the five first layers utilizing a rectified linear activation function (ReLU) and the last one using a sigmoid activation function, to provide final labels. The number of neurons in the hidden layers is 40,40,40,32,32 and 1, respectively.

Binary Cr oss-Entropy Loss or Logarithmic Loss.

- Adam optimization algorithm with learning rate=1e-5.
- Model trained for 60 epochs. Results are presented in Table 5.

**Table 5:**
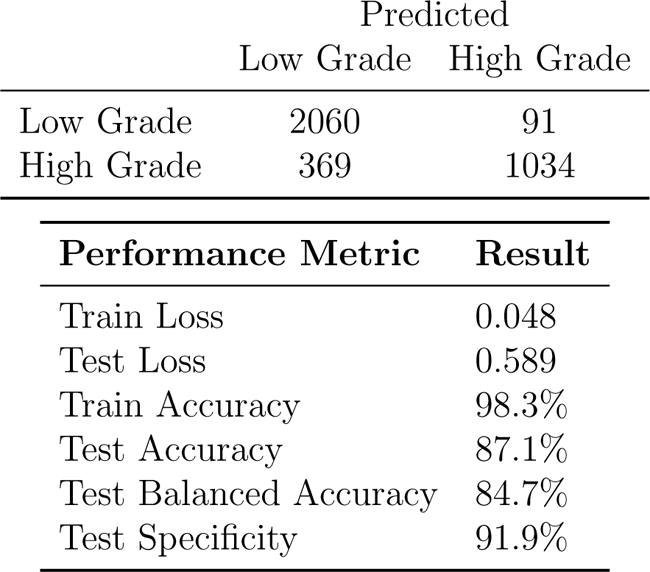
Low- versus high-grade distinction results with deep learning: confusion matrix and performance metrics.

#### 4.2.2 Normal versus abnormal distinction

The second classification task in this context was distinguishing between normal and abnormal histology findings using spectra data, motivated by the success in the low- versus high-grade classification problem. The most frequent label (abnormal) represents 32,793 spectra from the total of 34,870, corresponding to 94% of available data, an ever higher degree of data unbalance than the previous one. In this case, to provide better model guidance, a subset of train data was utilized as validation set. Several strategies were implemented to deal with this unbalance. The final model was selected according to the lowest false positive rate among models with similar test accuracy results. The neural network architecture and that achieved the best results had the following parameters:

- 5 hidden layers with the five first layers utilizing a rectified linear activation unit (ReLU) and the last one using a sigmoid activation function, to provide final classes. The number of neurons in the hidden layers was 40, 40, 32, 32 and 1, respectively. The first hidden layer also used a dropout rate of 50%.
- The weights were initialized with the logarithmic of the proportion between normal and abnormal spectra.
- Binary Cross-Entropy Loss or Logarithmic Loss.
- Adam optimization algorithm with learning rate=1e-5.
- Batch size of 2048.
- 300 total epochs.
- Early stopping protocol that interrupted training when precision in the validation set did not improve (mode=‘max’) for 10 epochs (patience=10).

The results are presented in Table 6.

**Table 6:**
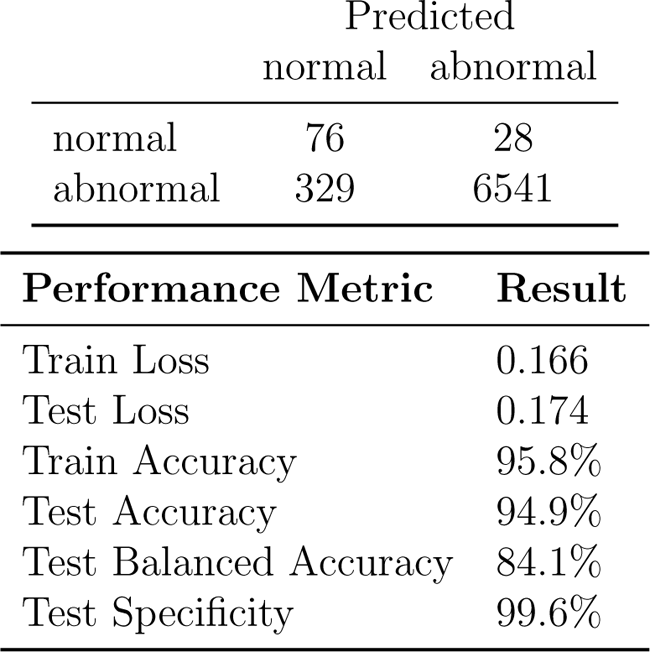
Normal versus abnormal distinction with deep learning: confusion matrix and performance metrics.

|  | Predicted |  |
| --- | --- | --- |
|  | normal | abnormal |
| normal | 76 | 28 |
| abnormal | 329 | 6541 |

| Performance Metric | Result |
| --- | --- |
| Train Loss | 0.166 |
| Test Loss | 0.174 |
| Train Accuracy | 95.8% |
| Test Accuracy | 94.9% |
| Test Balanced Accuracy | 84.1% |
| Test Specificity | 99.6% |

## 5 Discussion

### 5.1 Evaluation of Results and Implications

Machine Learning models in the normal versus abnormal tumor classification problem showed promising results. In this context, although Random Forest achieved the highest accuracy with 93.3%, SVM performed better in balanced accuracy and showcased a very good specificity, only 0.1% less than the best specificity, achieved by LASSO (95.8%).

The promising results in the aforementioned problem suggested that the same approach could be implemented in the second problem: low-grade and high-grade distinction. However, the results were not as satisfying, as can be seen in Table 4. It is important to note here that label proportion for the most common label here is 67.9% and some of the models did not even outperform the naive guess of all predictions following the most frequent label, while most of the others slight pass this threshold.

### 5.2 Deep Learning strategy

Given that distinction between high and low grade tumors did not perform well using models built with classical machine learning algorithms, this challenge was the first problem approached with deep learning techniques.

An ad-hoc method for deep learning training was implemented. The model was first evaluated by considering its accuracy and loss obtained in the training portion, followed by accuracy and loss obtained using a test dataset and its comparison with training results. If test accuracy results did not improve upon previous experiments or showed signs of overfitting — training results being much better than test results — model hyperparameters would be changed. This change was made either by returning to a previous iteration with better results or by evaluating a model history plot of loss and accuracy. The main relevant parameters in our deep learning problem were the learning rate in Adam optimizer and the number of epochs, alongside with changes in the number of hidden layers. In the optimization algorithm, smaller learning rates, varying in powers of ten, usually led to better accuracy results. We started with default parameters from the optimizer ([10]) and reduced from there. As expected, smaller learning rates led to longer training times. Our stop condition was based on lack of improvement in accuracy results. Given the high number of input parameters, a smaller learning rate provides us with the benefit of better mapping the possibilities of each derivative in the gradient.

Similarly, increases in the number of epochs were helpful to map through the possibilities of the data, especially because the number of patients is not high, even though the spectra count proved sufficient. They were usually evaluated through the history plots: if signs of noise were perceived in the results of any of the plots used or if there was a clear upward tendency in accuracy plots, the number of epochs was increased. Final plots for loss history in both classification problems can be seen in Figures 2 and 3. The resulting accuracy and losses were then compared with previous results and we checked for overfitting by comparing train and test results. When they were perceived as significantly different, model architecture would be changed.

**Figure 2:**
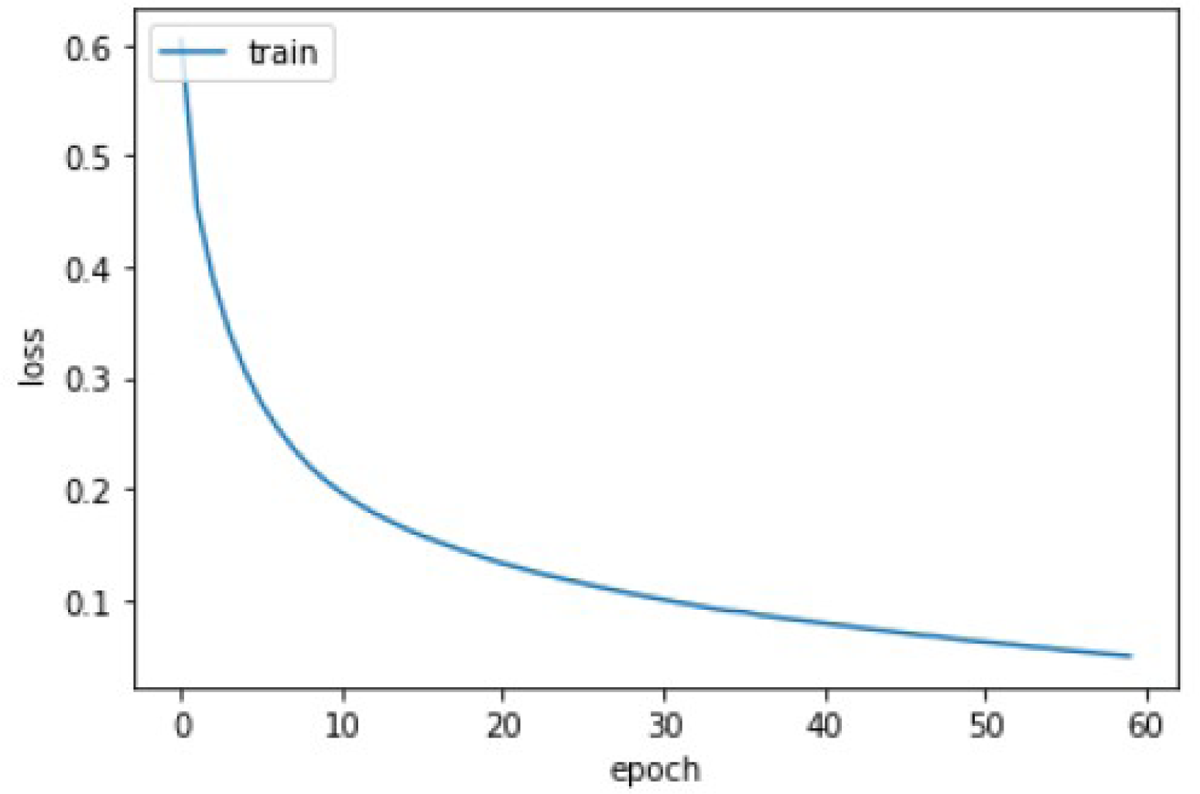
Train loss evolution plot for low versus high-grade classification.

**Figure 3:**
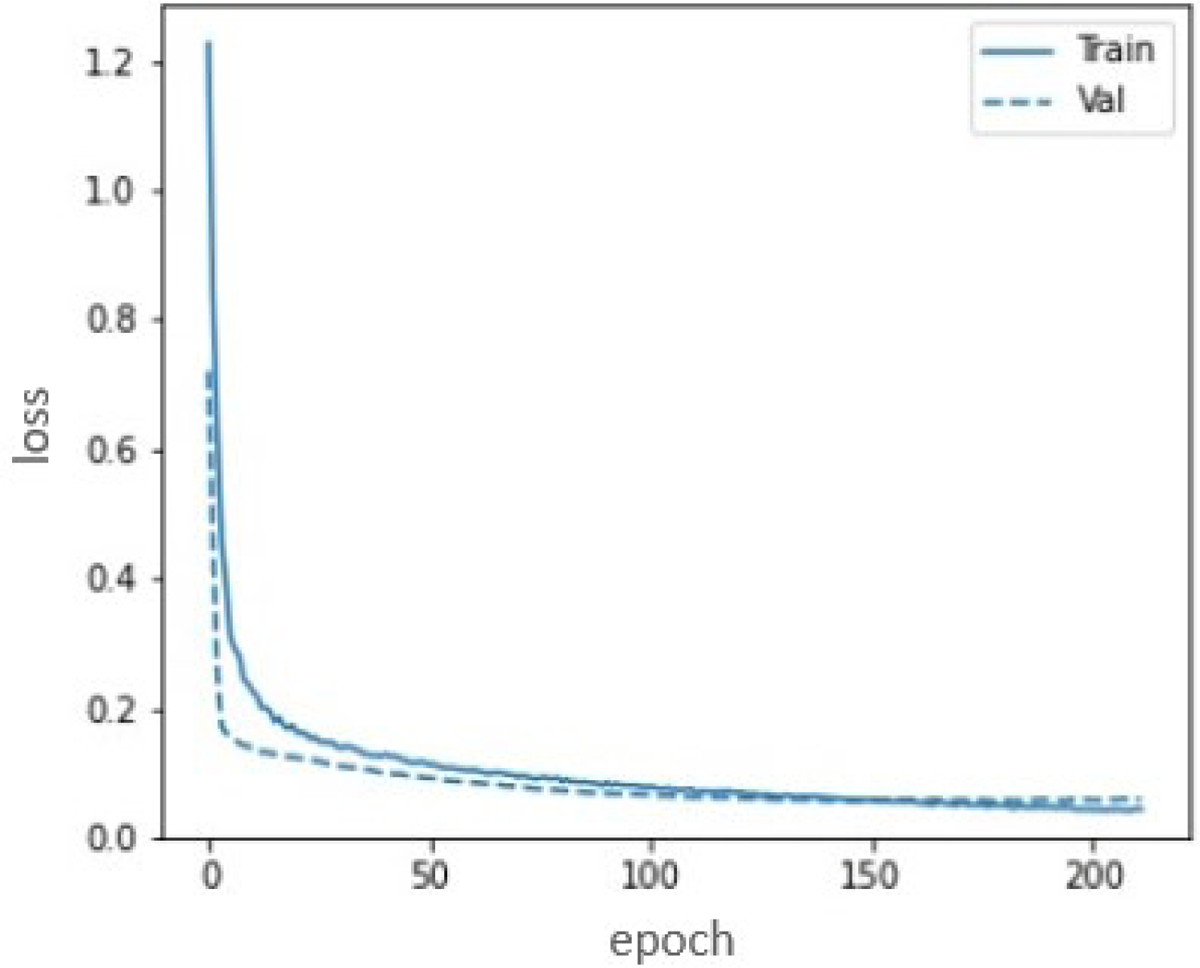
Train and validation loss evolution plot for normal versus abnormal classification. Due to the higher class imbalance in this problem, we opted to include a validation phase

Finally, the neural network architecture was evaluated in a similar manner. Whenever the change of parameters mentioned previously were not enough to achieve desired results, especially in comparison with machine learning results or previously achieved results, our approach was to change the neural network by adding or removing hidden layers.

In comparison with machine learning results, deep learning results were clearly superior, with test accuracy results being more than 20 percent points higher (Table 7). Also, similar results of accuracy and balanced accuracy were attained, which suggests a good overall performance in the distinction of every class separately.

**Table 7:** Summary of classification performance results for all models and tasks. Test Balanced Accuracy and Test Specificity are not reported for classical models in the low-grade vs. high-grade comparison because Test Accuracy values were already disappointing. Boldface marks the best results in each row.

| Normal vs. abnormal |  |  |  |  |  |
| --- | --- | --- | --- | --- | --- |
| Performance Metric | SVM | RF | MLP | LASSO | Deep |
| Test Accuracy | 88.6 | 93.3 | 92.3 | 82.3 | **94.9** |
| Test Balanced Accuracy | 71.5 | 70.9 | 67.9 | 66.1 | **84.1** |
| Test Specificity | 95.7 | 95.4 | 94.9 | 95.8 | **99.6** |
| Low- vs. High-grade |  |  |  |  |  |
| Performance Metric | SVM | RF | MLP | LASSO | Deep |
| Test Accuracy | 68.6 | 67.8 | 61.0 | 67.7 | **87.1** |
| Test Balanced Accuracy |  |  |  |  | 84.7 |
| Test Specificity |  |  |  |  | 91.9 |

### 5.3 Normal versus abnormal

The success in low versus high-grade problem lead to the desire to test a similar approach in normal versus abnormal problem. Test Accuracy results were initially very similar, but balanced accuracy was very low. The hypotheses was that those results were due to the fact that data imbalance was even more pronounced in this problem when compared with high versus low grade distinction. To tackle this problem, weight initialization was implemented, in order to manually suggest model imbalance. In order to further improve model performance in the case of high cardinality problems such as this, a dropout rate of 50% in the first hidden layer was also implemented. Training data was divided in batches to stabilize gradient search and an early stopping protocol was used to reduce training time. Scaling the data with Standard Scaler was tried but the performance decreased, so it was discontinued. Because of the fact that test accuracy was generally around 94%, the criteria to select the final model here was balanced accuracy and, finally, specificity, given the fact that a misdiagnosis of a abnormal tissue as normal could be more harmful in the context of an ongoing surgery for tumor removal procedure. The final performance history graphs for this problem are presented in Figure 3.

The final deep learning model outperformed machine learning models in all selected metrics, with a similar accuracy, but superior balanced accuracy and specificity, as presented in Table 7.

The emerging of high-throughput omics technologies and rapid advances in our understanding of the molecular basis of central nervous system tumors unveiled clinically relevant molecular classes and subgroups, pushing a new classification system by WHO in 2021 ([12]). While this updating is key to refine patient treatment options and improve quality of life, the next step towards precision oncology may be to implement molecular analysis techniques intrasurgically. This will leverage the concept of molecular resection margin, capitalizing on the new molecular classification potential and, ultimately leading to the development of subgroup specific surgical treatments.

### 5.4 Lessons learned

- Since one patient can have multiple spectra, they should be accordingly separated in order to avoid having the patient simultaneously in train and test sets. This can inflate accuracy results.
- Since the problem has significant heterogeneity, class unbalance should be taken into account when training the models.

## 6 Conclusions and Future Work

Our main conclusion is that it is worth using deep learning techniques for the classification problem of pediatric brain tissue into normal or abnormal, and into lowor high-grade, for malignant samples. More powerful computers are needed for training, but inference is fast even in modest computers, making it suitable for real-time applications such as during surgery.

In the future, we intend to add more data and redo the experiments, to increase confidence in the results, and perhaps extend the prediction to encompass all pathology classes. The inclusion of more sophisticated preprocessing steps, such as peak-to-peak filtering of spectra, is also worthwhile to try.

## Author contributions

TMSB: deep learning implementation, original draft preparation. MSD: classical method implementation. FC: MLP programmimg. DR: data organization. ALS: mass spectra collection, biological expertise. IAC: pathological classification. AMP: mass spectra collection, chemical expertise. HP: machine learning expertise. JM: conceptualization, writing, and implementation. All authors reviewed and approved the final version.

## Acknowledgments

We thank Drs Silvia Regina Brandalise and Pedro Otavio de Campos Lima for enlightening discussions about the clinical aspects of the project.

Funding: Brazilian Ministry of Health, by means of PRONON grant 25000.069610/2015-43 to Dr Izilda Cardinalli, as part of the National Program for Oncological Attention (PRONON), and by the Sao Paulo Research Foundation (FAPESP), by means of research grant 2018/00031-7 to Joao Meidanis.

